# Widespread separation of the polypyrimidine tract from 3’ AG by G tracts in association with alternative exons in metazoa and plants

**DOI:** 10.1101/363804

**Authors:** Hai Nguyen, Jiuyong Xie

## Abstract

At the end of introns, the polypyrimidine tract (Py) is often close to the 3’ AG in a consensus (Y)_20_NC<u>AG</u>gt in humans. Interestingly, we have found that they could also be separated by purine-rich elements including G tracts in thousands of human genes. These regulatory elements between the Py and 3’AG (REPA) mainly regulate alternative 3’ splice sites (3’SS) and intron retention. Here we show their widespread distribution and special properties across kingdoms. The purine-rich 3’SS are found in up to about 60% of the introns among more than 1000 species/lineages by whole genome analysis, and up to 18% of these introns contain the REPA G tracts in about 2.4 millions of 3’SS in total. In particular, they are significantly enriched over their 3’SS and genome backgrounds in metazoa and plants, and highly associated with alternative splicing of genes in diverse functional clusters. They are also highly enriched (3-6 folds) in the canonical as well as aberrantly used 3’ splice sites in cancer patients carrying mutations of the branch point factor SF3B1 or the 3’AG binding factor U2AF35. Moreover, the REPA G tract-harbouring 3’SS have significantly reduced occurrences of branch point (BP) motifs between the −24 and −4 positions, in particular absent from the −7 - −5 positions in several model organisms examined. The more distant branch points are associated with increased occurrences of alternative splicing in human and zebrafish. The branch points, REPA G tracts and associated 3’SS motifs appear to have emerged differentially in a phylum- or species-specific way during evolution. Thus, there is widespread separation of the Py and 3’AG by REPA G tracts, likely evolved among different species or branches of life. This special 3’SS arrangement contributes to the generation of diverse transcript or protein isoforms in biological functions or diseases through alternative or aberrant splicing.

## Introduction

Splice sites demarcate the boundaries between introns and exons for proper splicing of precursor RNA transcripts. Their sequences are constrained by a consensus but could be highly diverse among hundreds of species [1]. The diversity and flexibility may contribute to alternative pre-mRNA splicing as well as to the mutation effect in many diseases[2,3,4,5]. The majority of 3’ splice sites are comprised of the branch point, polypyrimidine tract and 3’ AG dinucleotides with a consensus (Y)_20_NC<u>AG</u>gt in humans based on the whole genome data[1]. These motifs are recognized by splicing complexes/factors U2 snRNP, U2AF65 and U2AF35, respectively[6,7,8,9,10,11]. Mutations of SF3B1 of the U2 snRNP complex and U2AF35 cause aberrant 3’ splice site usage in leukemia, breast and lung cancers[12,13,14,15,16].

Unlike the 3’SS consensus sequence where the Py and 3’AG are adjacent to each other separated by only two nucleotides ‘NC’, we have found a group of intron ends where they are separated further apart by RNA elements[3,4,17,18]. The first identified of these elements is CA-rich, called Ca++/calmodulin-dependent protein kinase IV (CaMKIV)-responsive RNA elements (CaRRE) to inhibit 3’SS usage through hnRNP L/LL [17,19,20]. We have further identified purine-rich including GGG elements within this region [18]. The G tracts inhibit U2AF65 binding and 3’SS usage through hnRNP H/F contributing to the emergence of novel alternative exons[3,18]. Together, we call these regulatory elements between the Py and 3’ AG REPA [18]. The REPA G tract-containing human genes are significantly enriched in cancer[3,18].

Analysis of individual 3’SS indicates that the human REPA G tracts were mostly ‘inserted’ between the Py and 3’ AG in the ancestors of mammalian genes during evolution[3,18]; however, their genome-wide prevalence and relationship to the upstream branch point and alternative splicing among different species remain unclear. In this report, we examine their distribution in individual 3’SS among >1000 Ensembl-annotated species/lineages, association with alternative splicing, their diverse host genes including those with aberrant splice sites in cancer, and association with distant branch point motifs.

## Results

### 1. Distribution of REPA G tract-harbouring 3’ splice sites among more than 1000 eukaryotic species/lineages

We first calculated and identified 17,643,684 annotated purine-rich 3’SS (⩾5 purines between the −10 and −3 positions) of 1175 eukaryotic species/lineages in the Ensembl releases R38/R91 (Fig. 1, & S_Table_I for matrices of each species and sequences of each 3’SS), as in our previous reports [1,18]. The purine enrichment contrasts the Py-rich content of the average 3’SS matrices of the human or other genomes[1]. The highest percentage of purine-rich 3’SS is 61% in the genome of fungus *Edhazardia aedis*, which is A/T-rich (38% each). Even in species that are highly constrained at certain positions for Ts within this region[1,21], such as the T_-5_ and T_-8_ in *C. elegans* and *B. microti*, respectively, and to a less extent the T_-5_ in *B. bigemina*, we still identified hundreds or thousands of purine-rich 3’SS (Fig. 1). The purine-rich 3’SS are most abundant in fungi, comprising about 17% of all of the 3’SS among the 434 species (Fig. 2A, mean ± SEM), followed by protists and plants, 14% and 11%, respectively, and the least (4%) in metazoans (particularly vertebrates, 3.5%, and mammals, <3%).

**Fig. 1.**
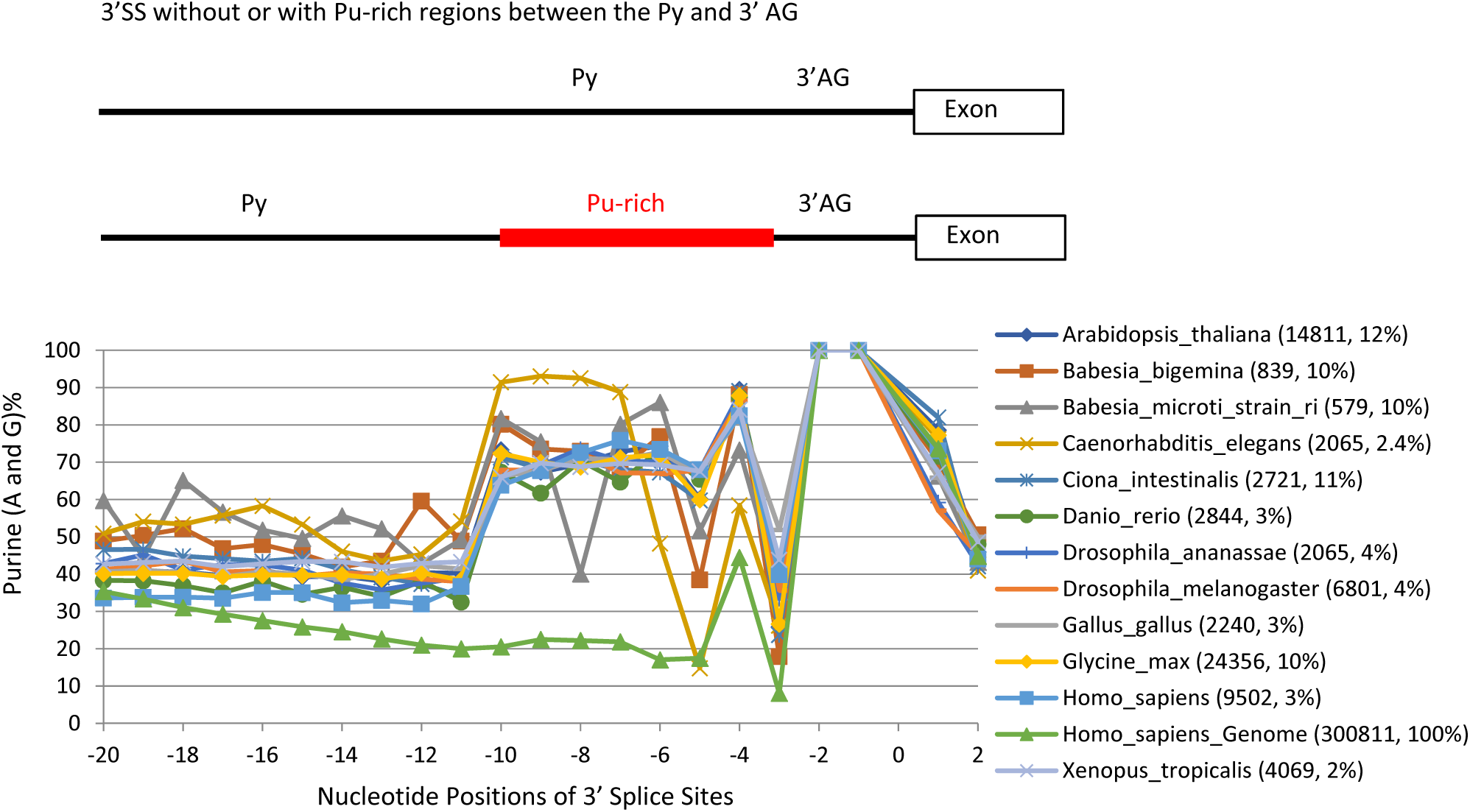
Percent distribution of purines (Pu) in the Pu-rich 3’SS between the Py and 3’ AG (−10 - −3) of a class of introns in genomes representatives of 1175 species/lineages. For convenience to trace the position-to-position changes in each species, their marked points of the purine nucleotide-percentages of each position are shown in straight line-linked curves. The distinct curve at the bottom between −20 and −3 is a control using the whole human genome. Bracketed next to the species names are the numbers of purine-rich 3’SS and their percentages of all 3’SS in the whole genome of each species. Position ‘0’ marks the intron-exon junction. Please see also S_Table I for the purine-rich 3’SS matrices of all the species/lineages, and S_file 1 for all the purine-rich 3’SS sequences.

**Fig. 2.**
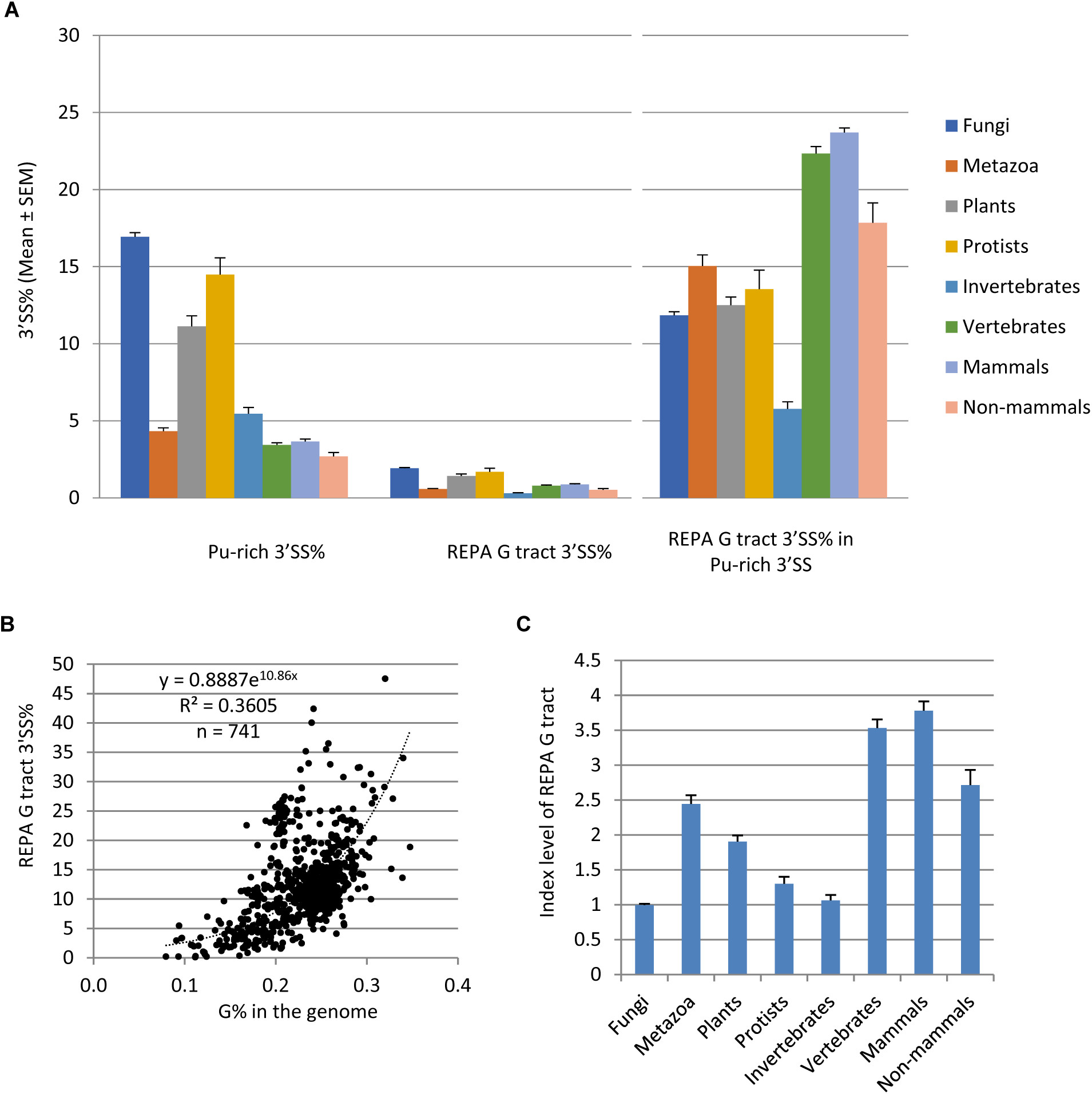
Distribution and enrichment of the REPA G tract motifs among different species/divisions. **A**. Percent distribution (Mean ± SEM) of the purine-rich 3’SS and REPA G tract 3’SS in the genomes of the four eukaryotic divisions, invertebrates and vertebrates of metazoa, mammals and non-mammals of vetebrates. Also shown is the percent distribution of REPA G tract 3’SS in the purine-rich 3’SS (Right panel). n = 434, 161, 52, 94, 71, 90, 69 and 21 species, for the 8 groups, respectively. Data from 741 unique species with >100 purine-rich 3’SS are included. **B**. Plot of the REPA G tract 3’SS% of the purine-rich 3’SS versus the guanine nucleotide G% of the genomes of the 741 species. The equation inside the graph is for the fitted trendline. **C**. REPA G tract enrichment index levels of the 6 groups (Mean ± SEM) after normalization to the purine-rich 3’SS% and genome G% of each species. For this, the REPA G tract 3’SS% in **B** was divided by the y values of the trendline corresponding to the G% of each species. The resulting mean value of fungal species is taken as 1.0 for comparison. Please see also S_Table II for the REPA G tract 3’SS matrices and S_Table III for the enrichment of REPA G tract 3’SS versus the purine-rich 3’SS and genome G% of all species/lineages, and S_file 1 for all the 3’SS sequences.

We identified 2,354,712 REPA G tracts, the most prominent of the 3’SS purine-rich motifs [3,18], between the −15 and −3 positions among 1,031 species/lineages (Fig. 2A, S_Table II and S_File 1 for matrices and sequences), ~13% of all the purine-rich 3’SS identified. Their percent distribution among the eukaryotic groups is similar to the purine-rich 3’SS except that vertebrates (mammals in particular) have higher level than invertebrates (Fig. 2A). However, relative to the purine-rich 3’SS, REPA G tract-harbouring 3’SS are most enriched (15%) in metazoans and the least (12%) in fungi among the four divisions. Of the metazoans, vertebrates have the highest level, ~22% (24% for mammals, 18% nonmammals), consistent with a previous genome-wide observation[22], while invertebrates have only ~6%. Upon further normalization to the G% in the genomes (Fig. 2C), the metazoa (vertebrates and mammals in particular) and plants are most enriched of the REPA G tracts (Fig. 2D). Therefore, separation of the Py and 3’AG by REPA G tracts is widespread and most enriched in metazoa and plants.

### 2. The REPA G tract-harbouring 3’SS are highly associated with immediate downstream alternative exons of metazoan and plant genes of diverse functions

The REPA G tracts regulate the alternative splicing of a large group of exons in human genes involved in cancer and cell cycle [3,18]. To determine if they are also associated with alternative exons among the other species, we examined all exons immediate downstream of the REPA G tracts for their constitutive or non-constitutive usage in the Ensembl database. Exons with both coordinates present in all transcripts in the database are taken as constitutive ones; or else, non-constitutive or alternative exons.

Compared to all exons in the transcriptome of each species, exons downstream of the REPA G tracts are significantly enriched with non-constitutive ones in 198 species/lineages (*p*<0.05, Fig. 3 & S_Table IV), and highly significant (*p*<0.001) in 123 lineages of 106 unique species. Of the latter, 73% are metazoan, 25% plant and less than 1% protist species (Fig. 3A). Of the metazoans, 89% are vertebrates and 11% are invertebrates. These comprise 48% of metazoan, 73% of vertebrates (74% of mammals and 71% of non-mammals) in particular, 52% of plant and 1% of protist species that contain the REPA G tract-harbouring 3’SS (Fig. 3B). Of the vertebrates, the primates night monkey *Aotus Nancymaae* and chimpanzee *Pan troglodytes* are among the top ones by enrichment folds and *p* values (Table I). Of the plants, the common wheat *Triticum aestivum* and cotton *Gossypium raimondii* are among the top ones. There are no fungi among the 106 species. Overall, the percentages of such AS-enriched species in each division or group is well correlated with the enrichment folds of the REPA G tract-harbouring 3’SS (Fig. 3B, R^2^=0.9162). Therefore, the REPA G tract-harbouring 3’SS are associated with alternative exons in more than a hundred species of mainly metazoans (vertebrates in particular) and plants.

**Fig. 3.**
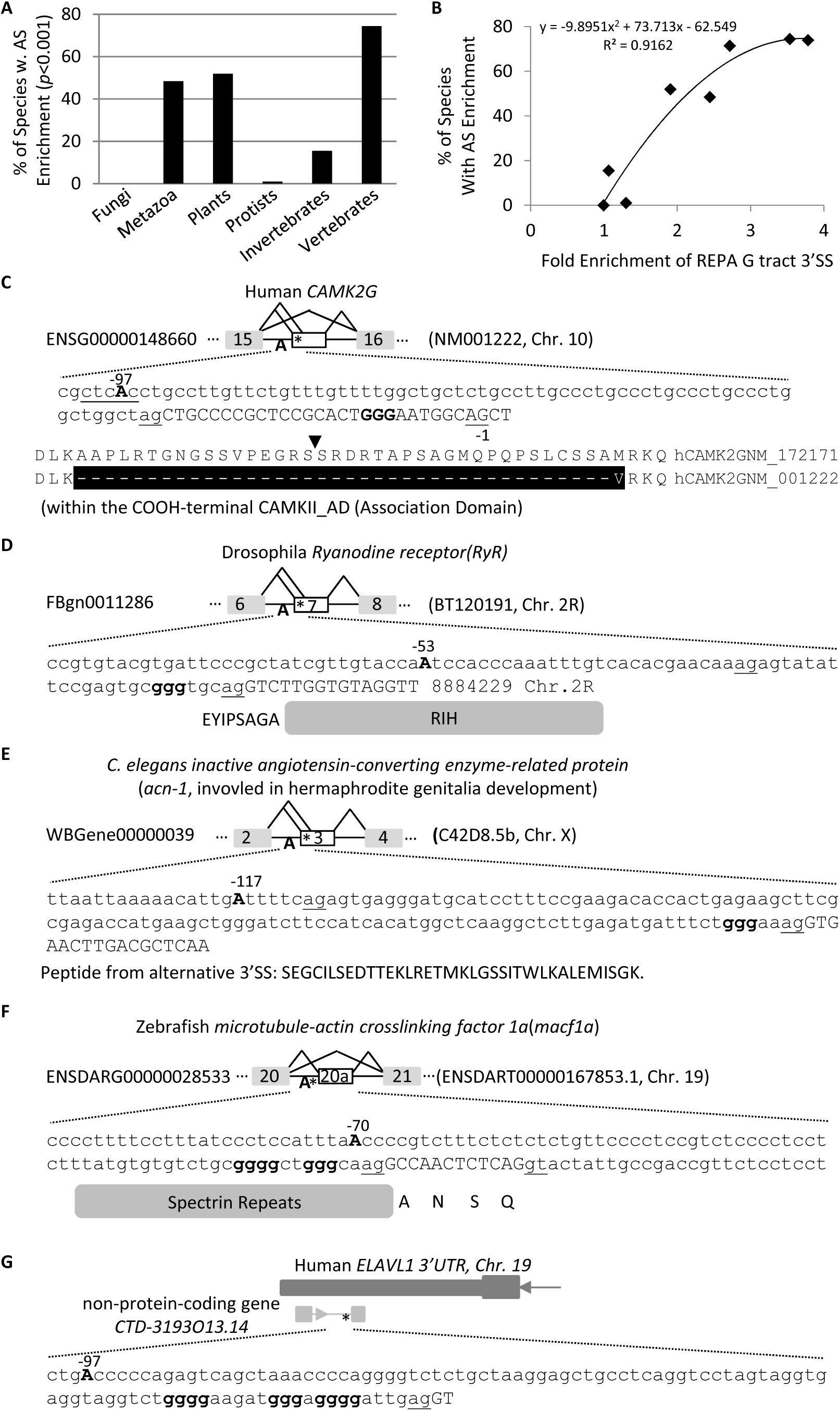
Enrichment of alternative exons downstrem of the REPA G tract 3’SS among 198 species/lineages. **A**. Percentages of species that show highly significant enrichment (p<0.001) of altenative exons downstream of the REPA G tract 3’SS in each division/group (n = 123 unique species in total). **B**. Correlation between the species percentages and the fold enrichment of REPA G tract 3’SS (over fungal species) in each division/group. **C**. Representative examples of 3’SS and alternative exons of protein-coding and non-protein-coding genes of several species. Uppercases are exon and lowercases intron nucleotides. ^∗^: GGG element. The 3’AG is underlined. The branch point Adenosine A is in bold and uppercase, and position is relative to the intron end G nucleotide (−1).

**Table I.**
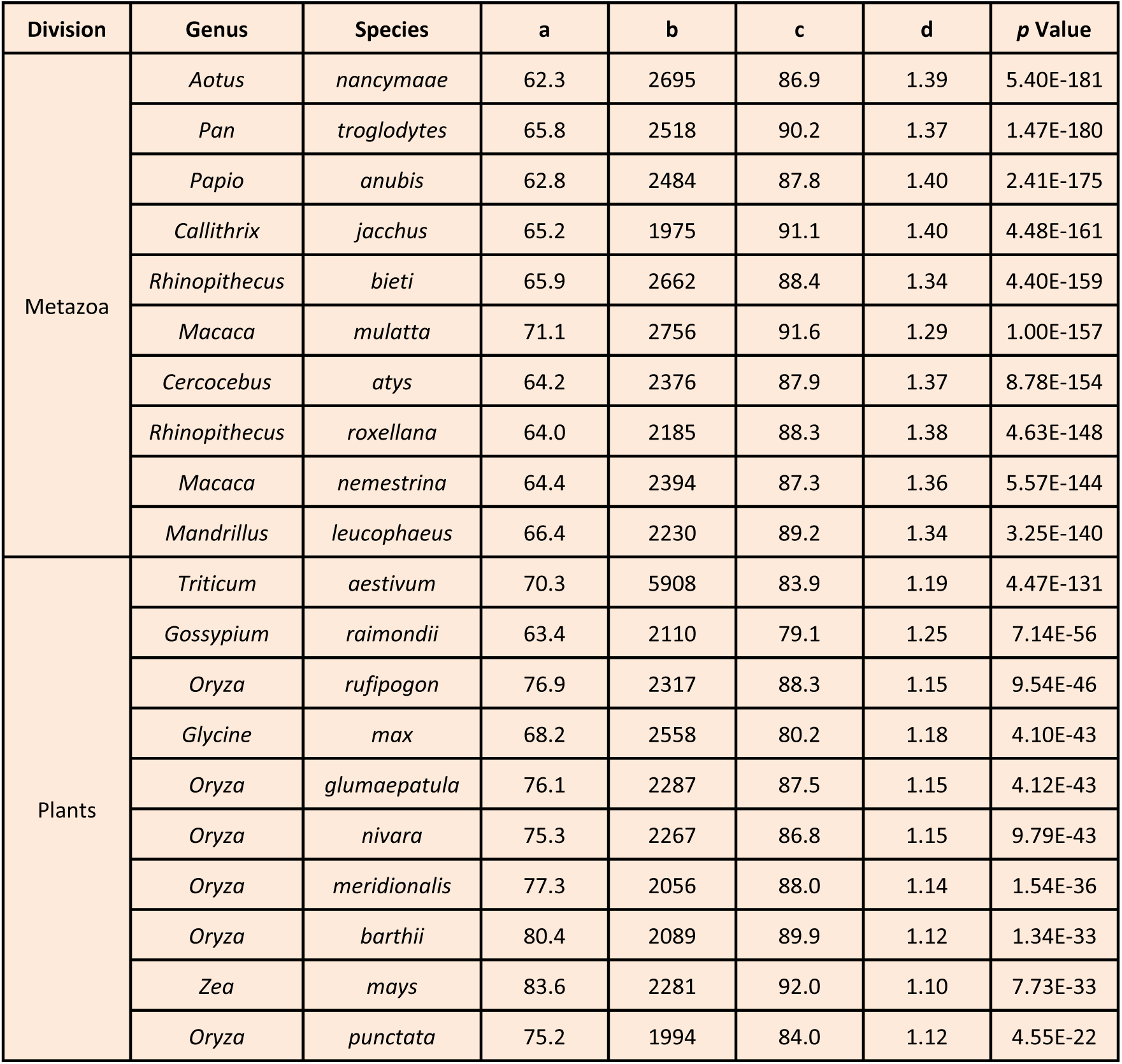
Top 10 species most significantly enriched of alternative exons downstream of REPA G tract 3’SS in metazoa and plants. **a**. percentage of non-constitutive exons in the transcriptome/genome of each species, **b**. total number of REPA G tract 3’SS, **c**. percentages of non constitutive exons downstream of **b**, **d**.Fold enrichment of **c** over **a.**

| Division | Genus | Species | a | b | c | d | p Value |
| --- | --- | --- | --- | --- | --- | --- | --- |
| Metazoa | *Aotus* | *nancymae* | 62.3 | 2695 | 86.9 | 1.39 | 5.40E-181 |
|  | *Pan* | *trogodytes* | 65.8 | 2518 | 90.2 | 1.37 | 1.47E-180 |
|  | *Papio* | *anubis* | 62.8 | 2484 | 87.8 | 1.40 | 2.41E-175 |
|  | *Callithrix* | *jacchus* | 65.2 | 1975 | 91.1 | 1.40 | 4.48E-161 |
|  | *Rhinopithecus* | *bieti* | 65.9 | 2662 | 88.4 | 1.34 | 4.40E-159 |
|  | *Macaca* | *mulatta* | 71.1 | 2756 | 91.6 | 1.29 | 1.00E-157 |
|  | *Cercocebus* | *atys* | 64.2 | 2376 | 87.9 | 1.37 | 8.78E-154 |
|  | *Rhinopithecus* | *roxellana* | 64.0 | 2185 | 88.3 | 1.38 | 4.63E-148 |
|  | *Macaca* | *nemestrina* | 64.4 | 2394 | 87.3 | 1.36 | 5.57E-144 |
|  | *Mandrillus* | *leucophaeus* | 66.4 | 2230 | 89.2 | 1.34 | 3.25E-140 |
| Plants | *Triticum* | *aestivum* | 70.3 | 5908 | 83.9 | 1.19 | 4.47E-131 |
|  | *Gossypium* | *raimondii* | 63.4 | 2110 | 79.1 | 1.25 | 7.14E-56 |
|  | *Oryza* | *rufipogon* | 76.9 | 2317 | 88.3 | 1.15 | 9.54E-46 |
|  | *Glycine* | *max* | 68.2 | 2558 | 80.2 | 1.18 | 4.10E-43 |
|  | *Oryza* | *glumaepatula* | 76.1 | 2287 | 87.5 | 1.15 | 4.12E-43 |
|  | *Oryza* | *nivara* | 75.3 | 2267 | 86.8 | 1.15 | 9.79E-43 |
|  | *Oryza* | *meridionalis* | 77.3 | 2056 | 88.0 | 1.14 | 1.54E-36 |
|  | *Oryza* | *barthii* | 80.4 | 2089 | 89.9 | 1.12 | 1.34E-33 |
|  | *Zea* | *mays* | 83.6 | 2281 | 92.0 | 1.10 | 7.73E-33 |
|  | *Oryza* | *punctata* | 75.2 | 1994 | 84.0 | 1.12 | 4.55E-22 |

We then examined the functional clustering of the host genes of representative metazoan and plant species using DAVID functional clustering analysis[23]. The clusters have common as well as highly specific ones among these diverse species (Table II). For example, the nucleotide- or ATP-binding cluster is found in *C. briggasae*, *Gorilla gallus*, *Homo sapiens* and *Pan troglodytes*, and the cluster calcium in drosophila, rat and plant *A. thaliana*, while as the cluster plastid is found in *A. thaliana* only. The clustered functional proteins range from membrane receptors, cytosolic signaling kinases, cytoskeleton/transport proteins, Golgi complexes as well as nuclear DNA/RNA binding proteins (Fig. 3C-F). Non-protein-coding RNA transcripts are also found (Fig. 3G). The resulting splice variants change the protein sequences or non-protein coding RNAs. Therefore, the REPA G tracts are associated with diverse common as well as specific functions among different species across the animal and plant kingdoms.

**Table II.**
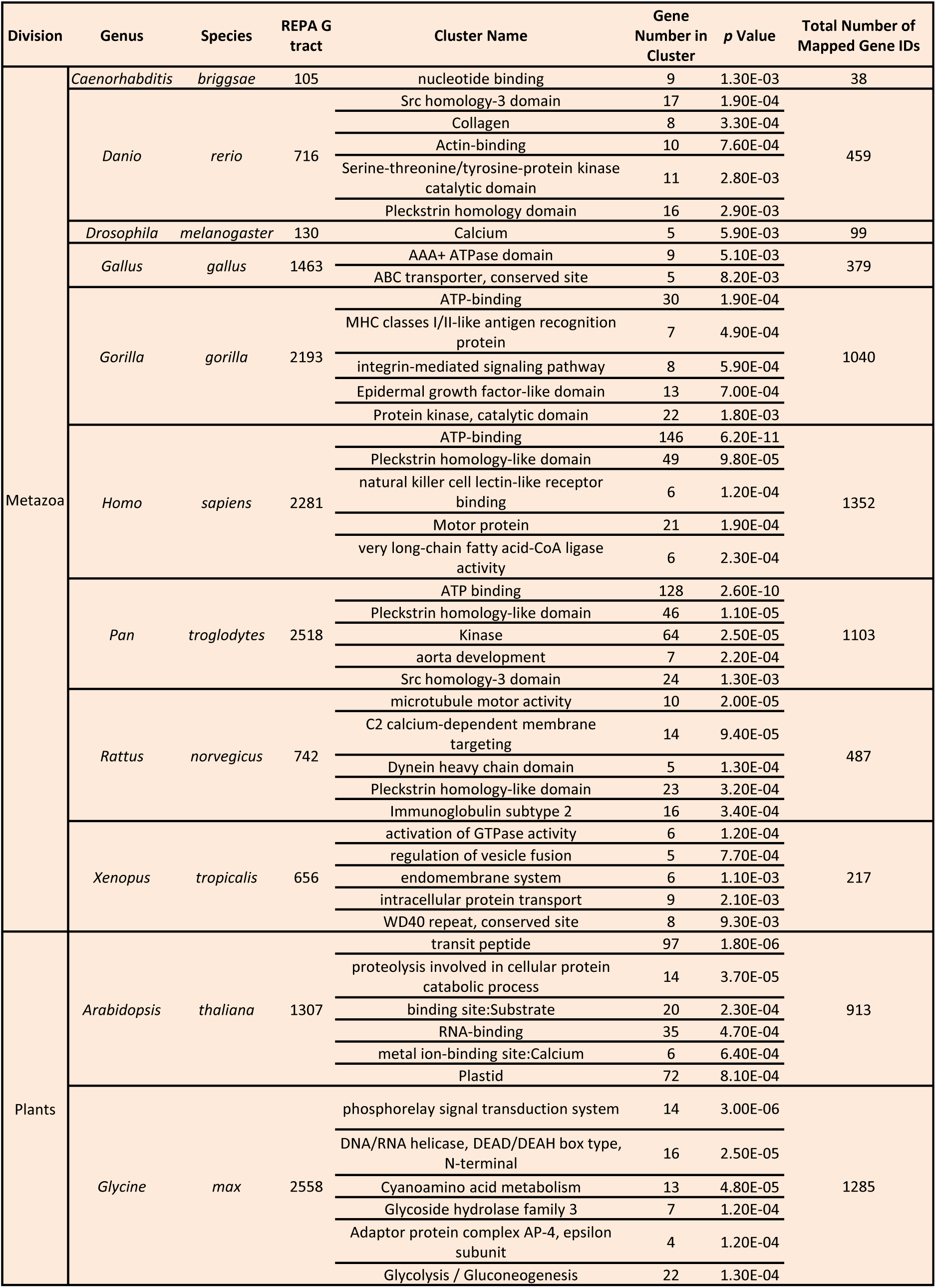
Most significantly clustered functions of genes containing the REPA G tract 3’SS and downstream alternative exons in representative species using DAVID functional clustering analysis.

### 3. The REPA G tracts are significantly enriched in the aberrant 3’SS that are caused by mutations of 3’SS factors in cancer patients

To assess the importance of these 3’SS in cell function or diseases, we examined their enrichment level in the well studied 3’SS factor mutant samples of human cancers. The REPA G tract-harbouring 3’SS comprise 0.76% of all 3’SS (or 24% of purine-rich 3’SS) in the human genome (Table III). They are relatively weaker splice sites that contribute to splicing inhibition through the bound hnRNP H/F[3,18]. We thus examined the REPA G tract-harbouring 3’SS usage when the 3’SS spliceosomal factor SF3B1 or U2AF35 are mutated in cancer or cultured cells [12,13,14,15,24]. Indeed, in SF3B1-mutated cancer patient samples[14], we identified 17 REPA G tract-harbouring 3’SS from 860 disrupted canonical 3’SS (2%, *p* = 2.68E-04), consistent with that these weak 3’SS are prone to be disrupted when the 3’SS factor is mutated. More interestingly, we identified even more (41) such G tracts from the paired aberrant 3’SS (4.77%, 6 folds, *p* = 4.21E-20, S_Table_Vb). Similar enrichment of REPA G tract-harbouring 3’SS is also found in a SF3B1m-expressing pre-B cell line Nalm6 samples[14] (Table III). Moreover, from the U2AF35m (S34F/Y) samples of lung carcinoma or acute myeloid leukemia [15], we identified 2.57% aberrant 3’SS that contain REPA G tracts (127 of 4947 aberrant 3’SS, S_Table_Va, *p* = 1.05E-31). Therefore, the REPA G tracts are not only enriched in the canonical but also even more so in the aberrant 3’SS upon SF3B1 or U2AF35 mutation in cancer patients or cultured cells.

**Table III.** Highly significant enrichment of REPA G tracts in the canonical or aberrantly used 3’SS in human cancers.

| Source of 3'SS | REPA G tract 3'SS | Total # of 3'SS | REPA G tract 3'SS% | p Value (vs Genome) |
| --- | --- | --- | --- | --- |
| Human Genome | 2,281 | 300811 | 0.76% |  |
| Canonical 3'SS_<br>SF3B1m_Cancer Tissues | 17 | 860 | 2% | 2.68E-04 |
| Aberrant 3'SS_<br>SF3B1m_Cancer Tissues | 41 | 860 | 4.77% | 4.21E-20 |
| Canonical_SF3B1m_Nalm6 | 8 | 591 | 1.4% | 4.54E-02 |
| Aberrant_SF3B1m_Nalm6 | 19 | 591 | 3.2% | 1.80E-07 |
| Aberrant 3'SS_<br>U2AF35m_Cancer Tissues/HeLa | 127 | 4947 | 2.57% | 1.05E-31 |

The aberrantly used alternative 3’SS and cassette exons in U2AF35m samples are enriched of the more prevalent CAG (65%) over TAG (27%) human intron ends [1,15,16,25]. Consistent with this observation, we found that the REPA G tract-harbouring 3’SS were enriched further by about 2 folds of CAG/TAG ratio over non-REPA G tract-harbouring aberrant 3’SS (Fig. 4A). The level is even higher for alternative 3’SS and cassette exons. Moreover, the most prevalent first nucleotide of exons is a G in the human genome (47%) [1]. The G_+1_ is slightly enriched among the 127 REPA G tract-harbouring 3’SS by about 1.2 folds (*p* < 0.02), and by 1.4 folds in the cassette exons (62%, *p* < 0.005), over non-REPA G tract-harbouring aberrant 3’SS. In contrast, the REPA G tract-harbouring 3’SS of the SF3B1m samples do not have as much enrichment or even reduction over the non-REPA G tracts, either aberrant or the corresponding canonical 3’SS. The G_+1_ increases the strength of the 3’SS in comparison with A_+1_ as measured by the MaxENT entropy scores[26] (Fig. 4B). Together, the aberrant REPA G tract-harbouring 3’SS are enriched of the more prevalent C_-3_ and G_+1_ over the non-REPA G tract-harbouring 3’SS in the U2AF35, but not SF3B1, mutants, particularly for the alternative 3’SS and cassette exons.

**Fig. 4.**
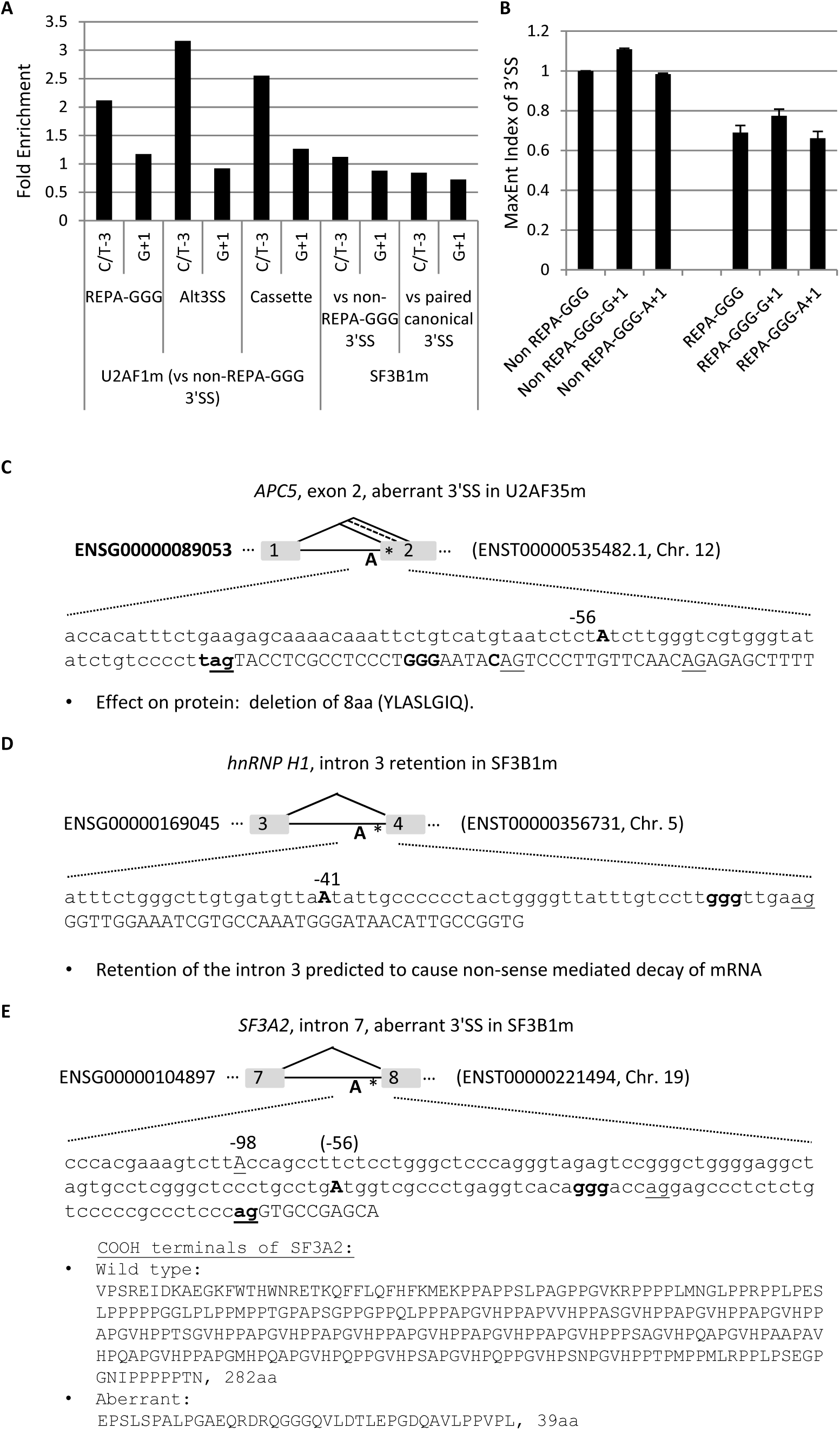
Enrichment of REPA G tract 3’SS in the aberrantly used 3’SS of U2AF35 or SF3B1 mutant human cancer samples. **A**. Fold enrichment of 3’ (C/T)_-3_ and G_+1_ over the non-REPA G tract aberrant 3’SS of the same mutant cancer samples. **B**. MaxENT scores of the different 3’SS. **C-E**. three examples of aberrantly used 3’SS in the U2AF35 or SF3B1 mutant cancer samples.

Three examples of such aberrant splicing of cell cycle regulators or of splicing factors related to the G tract or 3’SS control are in Fig. 4C-E: the *APC5* exon 2, *hnRNP H1* exon 4, and the *SF3A2* exon 8. In the *APC5*, a REPA G tracts aberrant 3’SS is used in U2AF35m samples, replacing the two flanking alternative 3’SS, resulting in a 24nt in-frame deletion of 8aa peptide YLASLGIQ. In *hnRNP H1*, the intron 3 was retained in *SF3B1m*, resulting in early termination of the ORF and expectedly nonsense-mediated decay of the transcript. In *SF3A2*, a 30nt-upstream AG (upAG) with a REPA G tracts in the intron was used to replace the wild type 3’AG in *SF3B1m*, causing a 28nt insertion and disruption of the open reading frame. The resulting protein is replaced of 282aa containing a PAT1 superfamily domain (Topoisomerase II-associated protein PAT1) by a shorter 39aa peptide. Therefore, usage of the REPA G tract-harbouring 3’SS in the *SF3B1* or *U2AF35* mutants affect cell cycle control genes, the splicing factors that target the GGG and other branch point factors.

Together, their enrichment in the 3’SS factor mutants of human cancer samples supports an important role of the REPA G tracts in cell function and diseases.

### 4. The REPA G tracts are associated with more distant branch point motifs

We have shown previously that the REPA G tracts are between the Py and 3’AG to weaken the 3’SS by their bound hnRNP H/F[3]. However, their relationship to the corresponding branch points remains unclear. A large group of the human branch points and consensus pentamer motifs (B-boxes) have been identified by lariat sequencing[27]. A number of the consensus motifs with predicted high affinity with U2 snRNA overlap well with experimentally verified ones[27]. We thus examined the position distribution of 5 of the top motifs (CTGAC,CTAAC,CTCAC,TTAAC and CTGAT) within the last 100nt of the introns among several representative species including human and vertebrate model organisms mouse, rat, zebrafish and a plant species (common wheat). The peak positions of the motif’s Adenosine are between −24 and −22 depending on the species, within the experimentally verified 90% of human BP regions between −19--37 with a median of −25 [27]. Interestingly, the BP motifs in the REPA G tract-harbouring 3’SS contain significantly less BP-A close to the 3’AG (−24 --4) than the control 3’SS or genome background in each species (Fig. 5A), in particular the BP-As are excluded from the −7 - −5 positions. This is accompanied by a significant increase between the −25/−24 and −97 positions (Fig. 5B). In humans, the BP-A peaks at −47 and −49 positions with significantly higher occurrences of alternative splicing (*p* < 0.005). An increase of alternative splicing was also seen for the zebrafish 3’SS with BP-As between −97 and −24 in comparison with that between −23 and −4 (p<0.05) but not in mouse and rat species. Therefore, the REPA G tracts are significantly associated with reduced BP motifs near the 3’AG and more distant BPs across the animal and plant kingdoms, and may contribute to alternative splicing in some species.

**Fig. 5.**
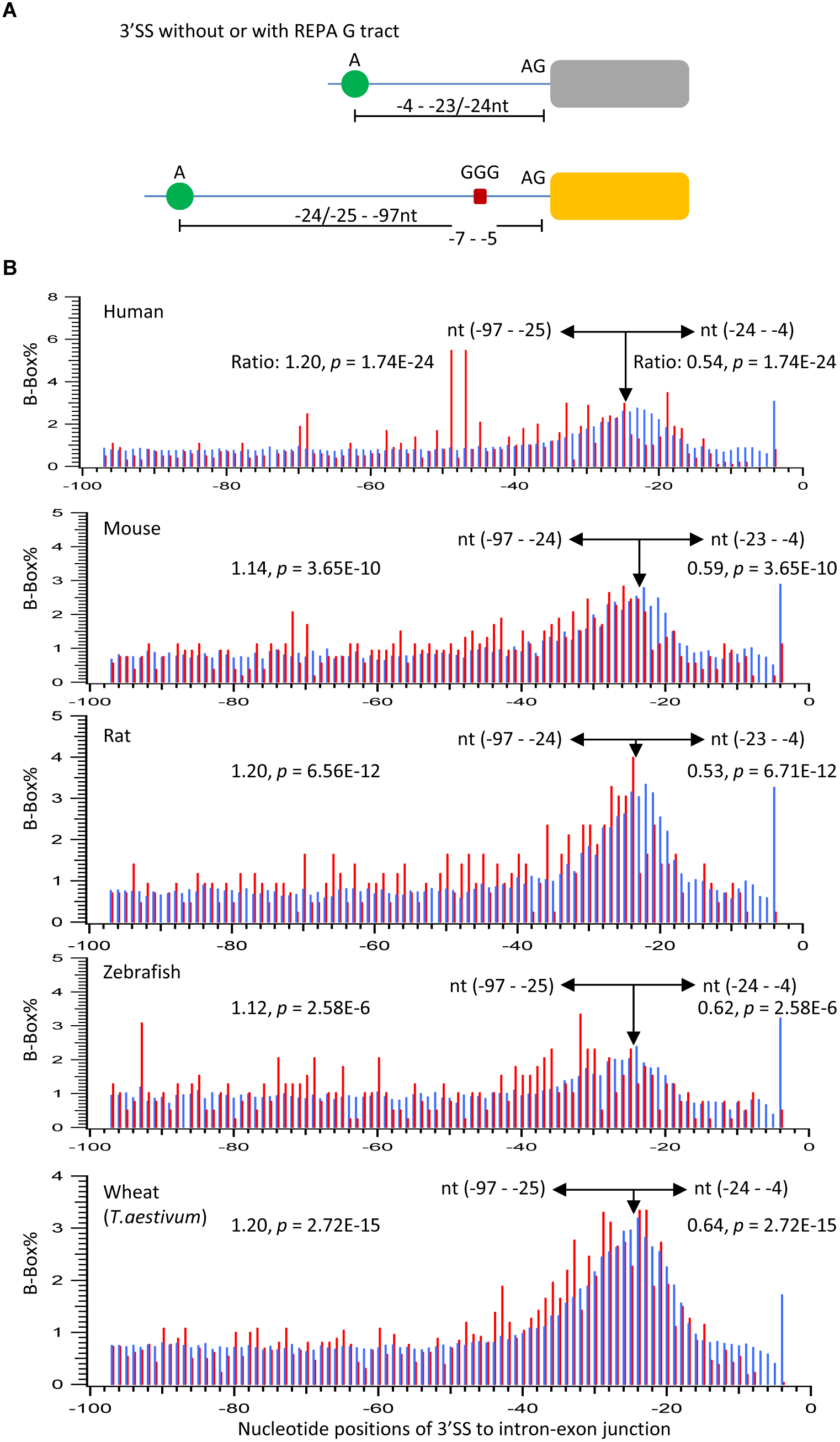
Reduction of branchpoint motifs (B-boxes) from the −24 - −4, and exclusion from the −7 - −4 regions in the REPA G tract 3’SS, accompanying their increase in the farther upstream regions from the 3’AG. **A**. Diagram of the major changes of the positions of branchpoint motifs in REPA G tract 3’SS. **B**. Percent distribution of the branchpoint motifs in the REPA G tract 3’SS (red) and the control 3’SS or the whole genome (blue).

Five examples of such REPA G tracts and distant BP motifs from human, drosophila, *C. elegans*, and zebrafish representing alternative 3’SS, cassette exon and intron retention events can be seen in Fig. 3C-G. The resulting variants from these alternative-splicing events changes or add peptides adjacent to protein domains or allow the generation of a non-protein-coding RNA transcript. The position of their predicted BP motif-As, according to the experimentally verified consensus motifs by Mercer et al[27], range from −53 to −117 upstream in the intron.

Further examination of these 3’SS motifs among the 100 vertebrate, 25 insect or 26 nematode species in the UCSC Genome Database indicated that the BP motif, upstream 3’AG (upAG), GGG and downstream 3’AG(dsAG) have highly variable levels of conservation (Fig. 6). They are conserved among most (79-88%) of the *CaMK2G* and *RyR* genes of vertebrate and insect species, respectively (Table IV). The two 3’AG motifs are also conserved among most (58-62%) of the nematode species. The zebrafish 3’SS motifs are mainly species-specific. Moreover, the human 3’SS of the antisense non-coding RNA at the 3’UTR of *ELAVL1* gene is mostly non-conserved except the downstream 3’AG among vertebrates. Together, these highly variable levels of species conservation of the REPA G tracts and associated 3’SS motifs suggest that the GGG motifs could have emerged either together or after the appearance of the nearby 3’SS signals in a species or phylum-specific way. Moreover, some species might have lost the motifs during evolution (in the cases of *CaMK2G* and *RyR*).

**Fig. 6.**
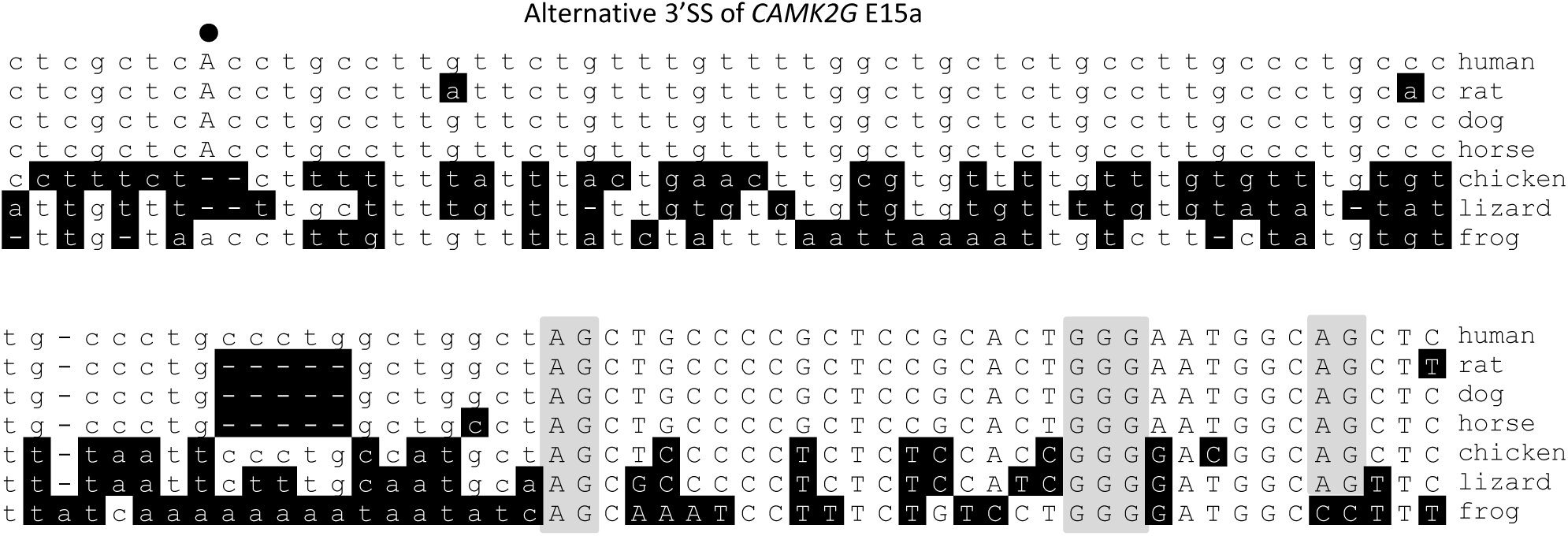
An example of the evolutionary changes of the REPA G tract and related 3’SS motifs, by Clustal alignment of the 3’SS of the *CaMK2G* exon 15a in Fig. 3C. Nucleotides different from the human gene is shaded in black. The 3’AGs and GGG are shaded in gray, and the potential branch point A in bold and uppercase.

**Table IV.** Percentages of conserved REPA G tracts and associated 3’SS motifs among the species of different groups. BP_A: the adenosine nucleotide of branchpoint, up: upstream, ds: downstream. H: humans, D: drosohphila, Ce: C. elegans, Z: zebrafish.

| Gene | BP_A | up3'AG | REPA G tract | ds3'AG | # of Species |
| --- | --- | --- | --- | --- | --- |
| *H_CAMK2G E15a* | 80% | 82% | 82% | 79% | 100 (vertebrates) |
| *D_RyR E7* | 80% | 88% | 88% | 88% | 25 (insects) |
| *Ce_acn-1 E3* | 8% | 62% | 8% | 58% | 26 (nematodes) |
| *Z_macf1a E20a* | 1% | 1% | 1% | 1% | 100 (vertebrates) |
| *H_ELAVL1 non-coding RNA, E2* | 14% | NA | 15% | 42% | 100 (vertebrates) |

## Discussion

In this study we have extended our previous finding about the REPA G tracts from mainly human to more than a thousand species across the eukaryotic kingdoms. They are as well highly associated with alternative splicing in more than 100 species of mainly vertebrates and plants, of genes with diverse functions and enriched in human cancers with 3’SS factor mutations. We also found their branch points are more distant from the 3’AG and associated with alternative splicing in some species.

### REPA G tract enrichment diversifies gene products in metazoa and plants through alternative splicing

The level of alternative splicing is higher in metazoa than that in fungi and protists [1,28,29]. Some plants also have relatively high levels (e.g. 40% of genes in maize *Zea mays* and 42% in *A. thaliana*)[30,31]. These high levels of alternative splicing require corresponding control RNA elements for proper splicing. There are 244,674 REPA G tracts in the metazoan and plant genomes (0.8% of all 3’SS, S_Table III, 1,064 ± 941, n = 230 species/lineages). Such widespread distribution together with their splicing inhibition effect suggests substantial contribution of these elements to the generation of splice variants in metazoa and plants.

We have shown that in humans these REPA G tract-containing genes are most significantly involved in cancer[3]. Here their widespread presence among different species goes far beyond cancer, to associate with a wide variety of gene functions including cell’s response to external environment (e.g. signalling, movement), as well as core DNA/RNA/glucose metabolism or highly specific plastid functions in plants (Table II).

In a step further from their association with cancer, we show that they are highly enriched in the aberrant 3’SS splice sites of cancers containing U2AF35 or SF3B1 mutations (Fig. 4 & Table III). Interestingly, the aberrant intron retention of hnRNP H1 is anticipated to cause NMD and reduced expression of the hnRNP H1[14] (Fig. 4), which could augment the usage of cryptic splice sites. Also the SF3A2 aberrant 3’SS usage results in protein truncation of a conserved domain, likely to weaken the U2 snRNP function as well, again increasing the usage of aberrant splice sites. Therefore, the SF3B1 mutation-induced splicing factor changes are expected to further enhance aberrant splicing in cancer patients. Together the REPA-G tracts represent a special class of RNA elements highly enriched in the aberrant splicing events in cancer.

### REPA G tract inhibition of 3’SS in aberrant splice sites and influence by distant branch points

It is not surprising that the REPA G tracts are enriched at the canonical 3’SS of mutant cancer samples since they are splicing silencers [3,18], which tend to make the host 3’SS skipped under suboptimal splicing conditions. What was surprising at first thought is the even more significant enrichment of these silencers at the aberrant 3’SS. However, their silencing effect is consistent with silencing the aberrant splice sites in SF3B1 or U2AF35 wild type cells. Their increased usage in mutant samples could be due to the better branch point consensus sequences of the aberrant 3’SS to match with U2 snRNA upon SF3B1 mutation[13,14], or by the more common CAG/G in U2AF35 mutants (Fig. 4)[1]. In these cases, the aberrant 3’SS are likely less influenced by the GGG.

We have shown previously that the REPA G tracts inhibits 3’SS usage by its *trans*-acting hnRNP H/F to interfere with U2AF65 binding [3,18], which is in a tight heterodimer with U2AF35[32]; thereby preventing their interaction with the Py and 3’AG, respectively[3,4]. Here they are also associated with more distant branch points from the 3’AG (Fig. 5). Even these are within 100nt in the intron end, not as distant as those found in other cases[33], they are still enriched of alternative exons in some species, consistent with a previous genome-wide observation on the effect of distant branch points [34]. Therefore, these GGG motifs likely regulate splicing with contribution from the more distant branch points as well.

A remaining question is what the functions are of such G tracts at the 3’SS of fungi and protists while they are not as enriched over the 3’SS and genome background as those in metazoa and plants. One possibility is that the corresponding 3’SS factors have evolved to be functionally compatible with the G tracts, as the 5’SS U1 snRNA or the LS2 protein in other cases [1,35], so that they are not as inhibitory of splicing as in humans[3,18]. Or they perhaps have other functions in RNA metabolism that remain to be identified.

G-quadruplexes close to the 5’ splice site have been reported to promote alternative splicing of the upstream exon [36]. However, they require at least four spaced repeats of GG, which are not present in the 2281 human REPA G tracts (S_file 1). Besides, the latters are 3’SS splicing silencers instead of enhancers. Therefore, the REPA G tracts apparently act distinctively on splicing from the quadruplex model.

In summary, the widespread separation of the Py and 3’AG by the presence of REPA G tracts across species of different kingdoms and their association with alternative exons indicate that these independently evolved regulatory elements and this unique class of introns contribute greatly to the transcriptome and proteome diversity through alternative splicing.

## Materials and Methods

### Genome data

The GenBank-format files of the genomes of all the species examined here were downloaded from the release 91 (mostly vertebrates) or Genome release 38 (invertebrates and others) of the Ensembl databases, of which the transcripts are based on experimental evidence[37].

### Analysis of constitutive vs non-constitutive exons

We compared the genome nucleotide position coordinates of all the exons of genes with more than one transcript in the Ensembl databases. Those with both coordinates present in all the transcripts are considered constitutive exons, or else, non-constitutive or alternative ones.

### Aberrant 3’SS in cancer patients or cell lines

These were obtained from the deposited sequences or related information from the published work by Darman et al. [14], or by Brooks et al. [15].

### Statistical Analysis

Hypergeometric test was used in the analysis of the density of REPA G tracts motifs and non-constitutive exons or positions of branch points.

## Acknowledgements

This work is supported by the Natural Sciences and Engineering Research Council of Canada (NSERC, RGPIN/6004-2016) and a Manitoba Research Chair fund to J.X.

